# ForestForward: visualizing and accessing integrated world forest data from the last fifty years

**DOI:** 10.1101/2024.12.05.626943

**Authors:** Eva Luz Tejada-Gutierrez, Jordi Mateo-Fornés, Francesc Solsona, Rui Alves

## Abstract

DATABASE URL: https://forestforward.udl.cat

Mitigating the effects of environmental exploitation on forests requires robust data analysis tools to inform sustainable management strategies and enhance ecosystem resilience. Access to extensive, integrated plant biodiversity data, spanning decades, is essential for this purpose. However, such data is often fragmented across diverse datasets with varying standards, posing two key challenges: first, integrating these datasets into a unified, well-structured data warehouse, and second, handling the vast volume of data using big data technologies to analyze and monitor the temporal evolution of ecosystems. To address these challenges, we developed and used an ETL (Extract, Transform, Load) protocol that curated and integrates 4,482 forestry datasets from around the world, dating back to the 18th century, into a 100GB data warehouse containing over 172 million records sourced from the GBIF (Global Biodiversity Information Facility) repository. We implemented Python scripts and a NoSQL MongoDB database to streamline and automate the ETL process, using the data warehouse to create the ForestForward web platform. ForestForward is a free, user-friendly application developed using the Django framework, which enables users to consult, download, and visualize the curated data. The platform allows users to explore data layers by year and observe the temporal evolution of ecosystems through visual representations.

## Introduction

The interconnectedness of human, environmental, and animal health, recognized in the One Health concept (1), underscores the need for sustainable management of ecosystems and biodiversity. Better environmental policies decrease the probability of zoonotic events (2), such as COVID19, and help mitigate other consequences of environmental degradation (3). These events are becoming more frequent because of human activities and climate change that are altering the planet’s ecosystems (4, 5). We need collaborative, interdisciplinary approaches to achieve optimal health for people, animals, and the environment (6). These approaches require collect and integrating high-quality data about the environment, animals, and humans. However, data sources are often too diverse and fragmented, posing considerable challenges to data integration (7). In addition, satellite technology is creating new data sources that will be crucial for an appropriate management of ecosystems (8, 9). Still, there are decades, and in some cases, centuries of information regarding the abundance and biodiversity of plants around the world that could help us understand how the green coverage of the planet evolved over large periods.

Many specialized databases contribute to the understanding of global flora. For example, the Global Root Traits (GRooT) (10) provides a complete database specializing in the characteristics of the plant’s roots, while the Global Naturalized Alien Flora (GloNAF) provides data on plant species that were introduced to regions outside their native range (11). The Global tree portal search offers a searchable database of tree species by scientific name and country (12), while World Flora Online (WFO) provides global taxonomic data, including images of species (13). As a final example Global Inventory of Floras and Traits (GIFT), adds a geographical component, allowing users to explore plant species and traits through a map-based interface (14). However, despite the wealth of data these and other platforms provide, they often remain isolated, and efforts to integrate multi-source data are limited.

Two major initiatives have been instrumental in collecting flora data globally, each with its own strengths and limitations. The Global Biodiversity Information Facility (GBIF) is an international platform that provides information from 1760 national, regional and local institutions including the European Environment Agency and the Atlas of Living Australia (ALA). GBIF collects data about biodiversity from around the world and promotes open data access, making biodiversity data freely available to researchers and providing informatics tools to facilitate biodiversity studies (15). However, the quality of the various data sets is highly uneven, which can hinder an integrated utilization and limit their usefulness. The other initiative, Global Forest Biodiversity Initiative (GFBI), imposes a stricter and uniform data format and provides controlled access to recent, high-quality forest biodiversity data (16). While the standardized approach of GFBI enhances the reliability of data for specific analyses, the restricted access to its datasets limits the potential for widespread use, especially by the broader research community.

Combining the strengths of open access data platforms like GBIF with the structured approach of GFBI can significantly improve the capacity for temporal analyses in biodiversity studies. By integrating diverse datasets into a unified framework, researchers can develop more accurate models to predict the effects of climate change and human activities on ecosystems. This, in turn, can support more effective conservation strategies and contribute to macroecology and biogeography studies, which require large-scale, spatiotemporal data on species distributions and ecosystem dynamics. (17, 18).

Here we focus on developing and using Extract, Transform, and Load (ETL) tools to integrate thousands of diverse plant abundance data sets from around the world, over large periods into a unified knowledge base, ensuring data consistency. We extract the datasets from GBIF and gather them into a unique database, accessible via the ForestForward web platform. ForestForward follows FAIR and Open Science principles, and enables researchers to visualize and download data in uniform format, dating back to the 17th century.

### Methodology

#### Data source and download

The data used in this work were downloaded from GBIF. The first download was done using a link sent by email after a request in the GBIF web page (GBIF.org, 2018). Data was originally downloaded on May 3, 2018. After that, we developed a Python bot that, every tenth day of the month, accesses and processes a csv file from GBIF with all new flora data from the previous month. New data sets are continuously downloaded and integrated. Supplementary Table S1 contains a list of all datasets integrated into ForestForward until September 2024.

#### Database technology

We chose MongoDB as the database technology for ForestForward. This No-SQL motor manages small and large volumes of data in a flexible and reliable way (19). Documents are in Javascript Object Notation (JSON), but saved in BSON format in MongoDB, which is a format that permits saving the fields inside the document as a key-value pair. We use the GEOJSON format (20) to save geographic data and perform queries related to location inside a polygon. We defined indexes in the collections to speed up the queries. MongoDB technology facilitates the integration of data from different sources because the data have different features. It also ensures scalability as the database grows larger (21).

#### Data integration and curation

We designed and implemented a structure that partially replicates the one reported by GFBI (Supplementary Figure S1) for the ForestForward database. The main difference with respect to GFBI’s database is that we do not store plant diameter and height, because this data was absent from the vast majority of integrated datasets. Figure 1 shows that ForestForward relies on a main knowledge base (Plantae), and an ancillary data collection (Grid). Plantae saves the individual records from all datasets in a uniform structure. It currently contains over 100GB of data. The most important fields of this collection are the taxon (species, genus) and geographical coordinates. We defined geographic coordinates (latitude and longitude), species, genus, and year of observation as key fields, in order to speed up searches and guarantee accurate representation on the map. Those fields uniformize the various datasets and facilitate any future integration with GFBI’s knowledge database format, which also contains them. Grid contains preprocessed information that allows ForestForward to work faster for standard tasks, by lumping data across periods of one year and by geographic location. The Grid collection has a summary of the amount of documents pre-processed and grouped by geography location, number of species and temporality of the observation.

**Figure 1.**
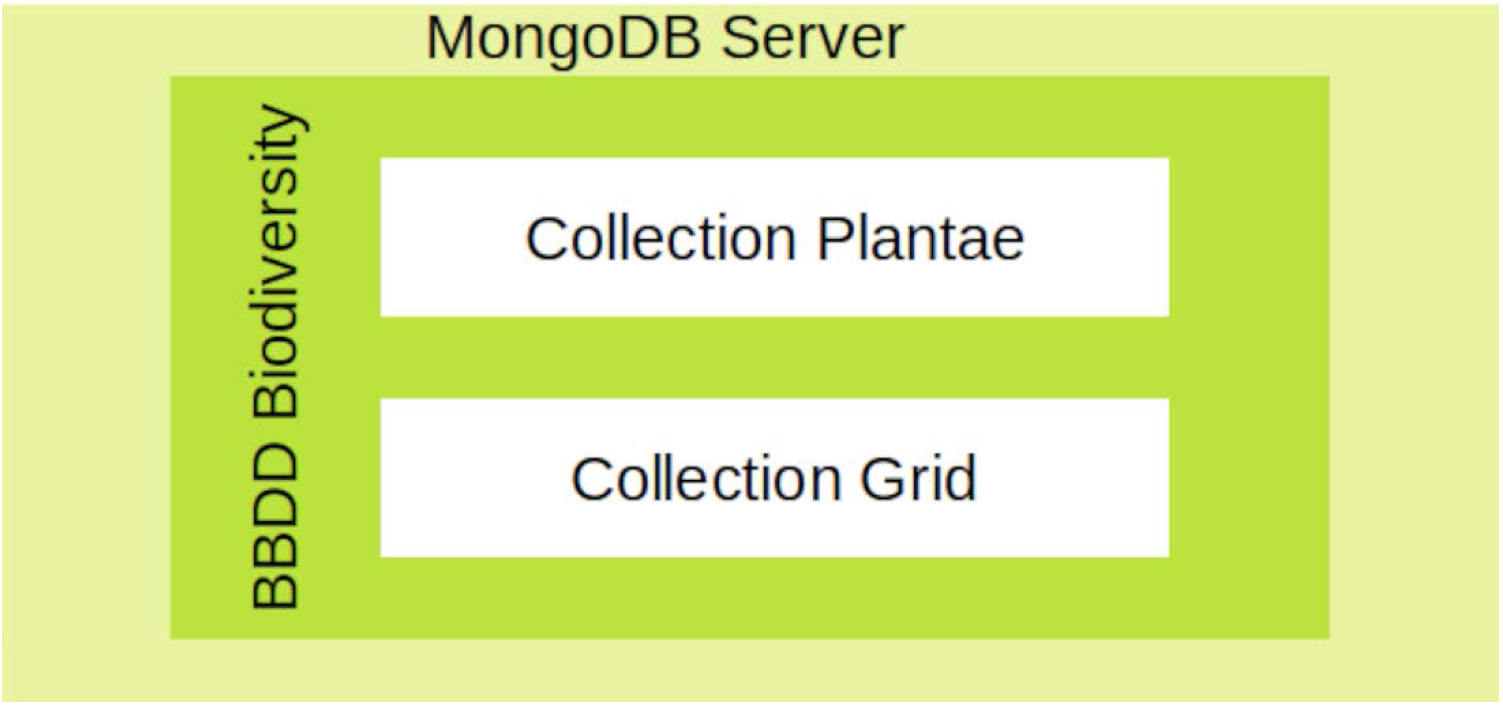
The database architecture designed for ForestForward. The main collection, Plantae, contains all records, while the ancillary collection, Grid, contains preprocessed documents to increase the speed of search.

Data curation is done through a semi-automatic process. On the one hand, the curation process looks for duplicate records in new data, before integrating it into the Plantae collection. This search is done by matching source id, latitude and longitude, species, genus, and year of observation. If duplicates are found, they are not included in the database. On the other hand, incomplete records that lack any of the key fields are stored in a temporary collection, to be revised after automated integration of complete records. During revision, the only records that are integrated into the main Plantae database are those whose only missing information is the species field. We note that these records are also matched to the Plantae collection in search of duplicate records. We discard the new, incomplete, entry if all its fields fully match any pre-existing record. We download new data every month and include it into the Plantae collection, after curation. The new data is also used to recalculate the information and update the Grid collection.

#### User interface and Database integration

ForestForward’s interface was designed to make the database available to the community and enable basic descriptive statistics of the data in a user-friendly manner. The application has a Client-Server architecture developed in the Python-based Django framework (22) that uses the Model-View-Controller (MVC) pattern (23). The Model layer is related to data, the View layer is related to the application interface, and the Controller layer manages the view requests, querying the model and response to the view. MVC is a design pattern that allows modular independence to the application. With MVC, web views can be modified without affecting the structure and model of the web application.

We achieve the connection between Django and the MongoDB database through the Mongo engine, which maps the data in the Model layer. We use the Folium JavaScript library (24) to represent the world map and overlay the plant data. Statistics and plots are created with the JavaScript library Char.js, which creates maps requiring less memory use, because plots are built during execution time of the query.

## Results

### Extract Transformation Load process

One of the most important processes is obtaining the data from the thousands of GBIF datasets, formatting them uniformly and integrating everything into our database. We automated these tasks by developing and applying a Python script, summarized in Figure 2.

**Figure 2.**
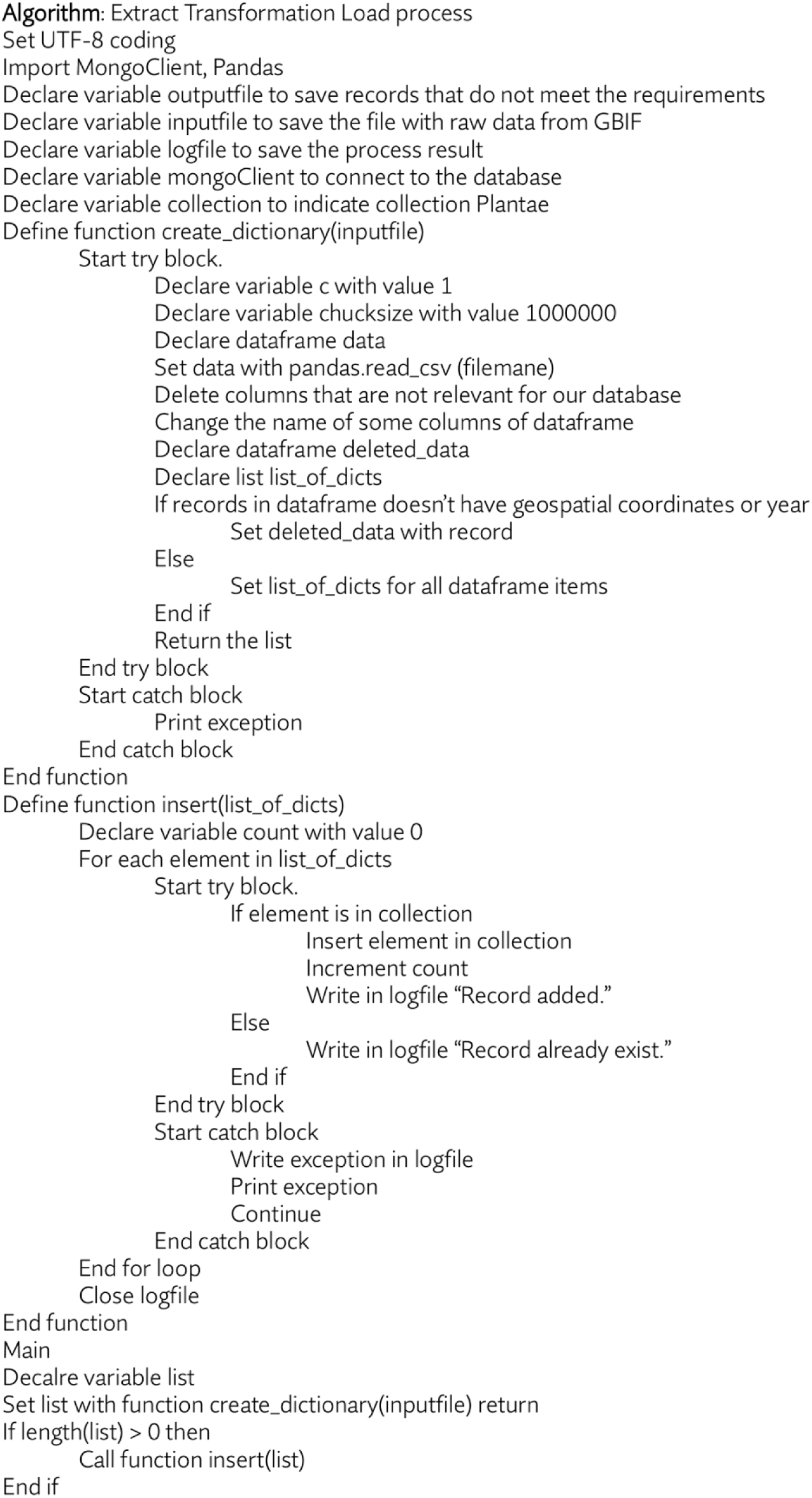
Pseudo code for the Python script that identifies, downloads, formats, and integrates new GBIF datasets into our database.

### Database design and implementation

Every document in the Plantae collection has twenty-seven fields. We identify each document with an objectID field called *_id*. Five database fields maintain traceability of the records to their original datasets: Gbifid, taxonkey, specieskey, datasetkey and occurrenceid. The first three are of type numeric, and the remaining two are of type string. The four fields Kingdom, phylum, species and scientificname are of type string and contain taxonomic information about the record.

There are nine fields containing geographic information. Countrycode is a string field with the ISO 3611 country code (25), locality is a string with the location indication. Decimallatitude, decimallongitude, coordinateuncertaintyinmeters, elevation and depth are numeric fields of coordinate values. Geoloc is a GEOJSON object of type point with the coordinate pair Latitude-Longitude. An array field called grid5 contains the pair Latitute-Longitude of the polygon to which the document belongs to in the Grid collection. There are three fields about precision of geographic information. Coordinateprecision, elevationaccuracy and depthaccuracy are numeric fields that record precision with respect to the coordinates, elevation or depth fields (See Supplementary Figure S1 for details).

Finally, there are four fields containing information about the date in which the record was taken: Eventdate, day, month and year. Eventdate is a string, while the last three fields are of type numeric. There is one additional field that is a flag to identify the reincorporated records after the quality review where species was replaced by genus.

Documents in the Grid collection have six fields of preprocessing data. The first field is for the identification of type ObjectID called _id. The second field is an array of two elements that contains the vertices of the polygon. The third field contains the preprocessed information about the year of observation. TotalRegistros and totalSpecies are numeric fields with the amount of records preprocessed in this square. Finally, the sixth field, specieList, is an array that has the list of species and the number of records per species in this polygon. This information is summarized in Supplementary Figure S1.

### Data description

From the more than 3500 original datasets, after curation and integration, 109 955 246 records were integrated into the ForestForward data warehouse. The curation process discarded 93 407 599 incomplete and 8 821 991 duplicated records, eliminating 49% of the raw data.

77 687 228 new documents were downloaded between 2018 and 2024. Of these, 62 139 537 were finally incorporated into the ForestForward database (26–34) (GBIF.org 2019, 2020, 2021, 2022, 2023, and 2024) (Figure 3). The percentage of discarded data in each update of the data warehouse is widely variable. In a few occasions no new data was discarded, while in others up to 68% of the new entries did not meet our defined quality standards for integration. Overall, 41.5% of all raw data was not integrated into the ForestForward data warehouse, as we can see in Figure 3. Currently, the database contains information about 307 898 species in 172 094 783 documents, each with their respective objectID. We note that these number change with each update.

**Figure 3.**
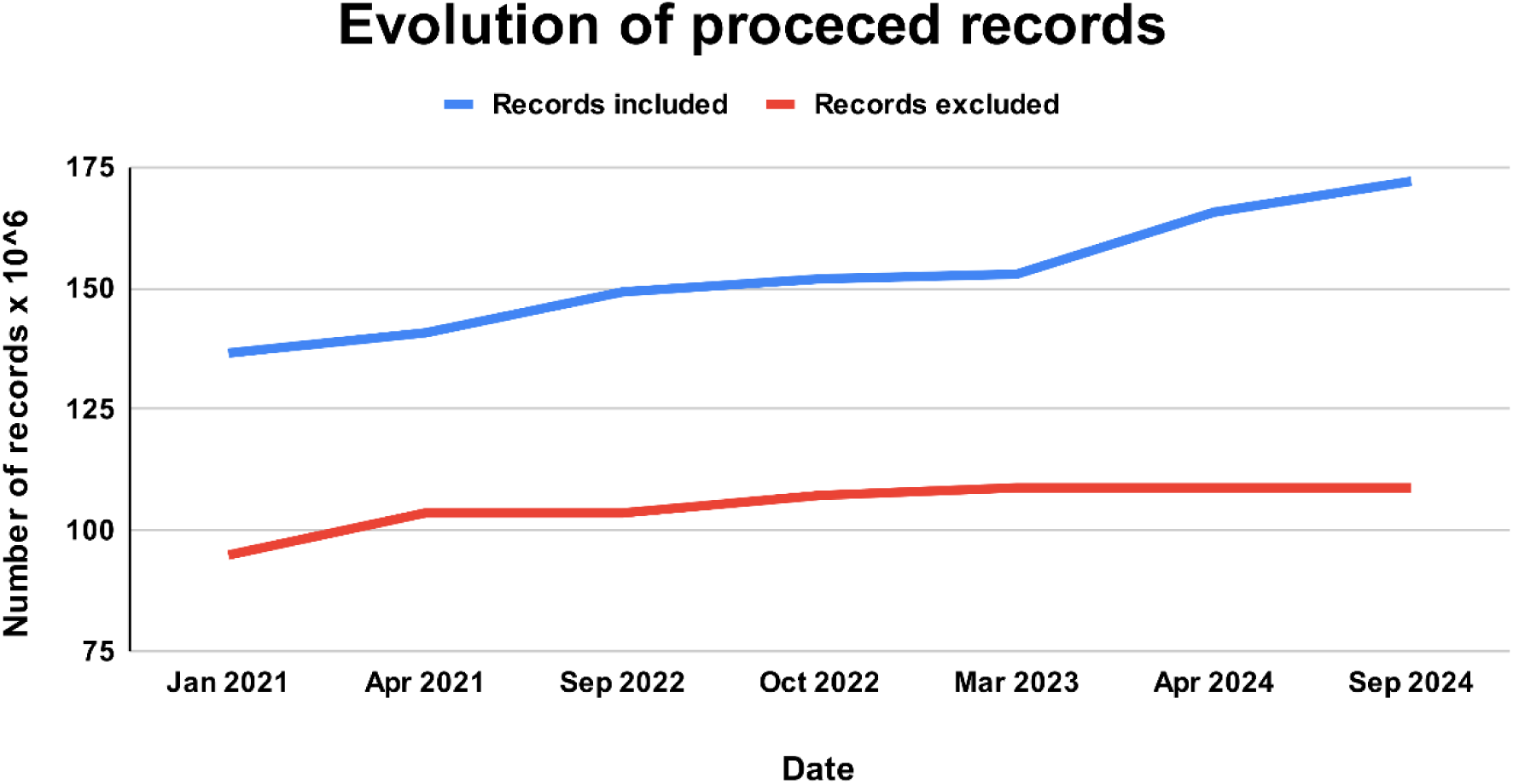
Evolution of records processed and integrated to ForestForward. Blue - Total number of curated records included in our database from GBIF. Red - Total number of records from the GBIF datasets that were excluded from our database due to duplication or low data quality.

### Interface and Functionality

ForestForward is available at https://forestforward.udl.cat. Figure 4A shows the entry screen for the web page. Users have two ways to query the data. First, they can go to the search page and filter the data by year, species, and/or country (Figure 4B). A search and download button in this view generates two downloadable files for each search. One file contains the filtered raw data in cvs (Comma Separated Values) format. This file contains species, year, country, latitude, longitude and datasetkey fields for each record. The datasetkey field traces the record to its original dataset. The other file contains a basic descriptive statistics summary for the number of records that were found in each year and country for the search.

**Figure 4.**
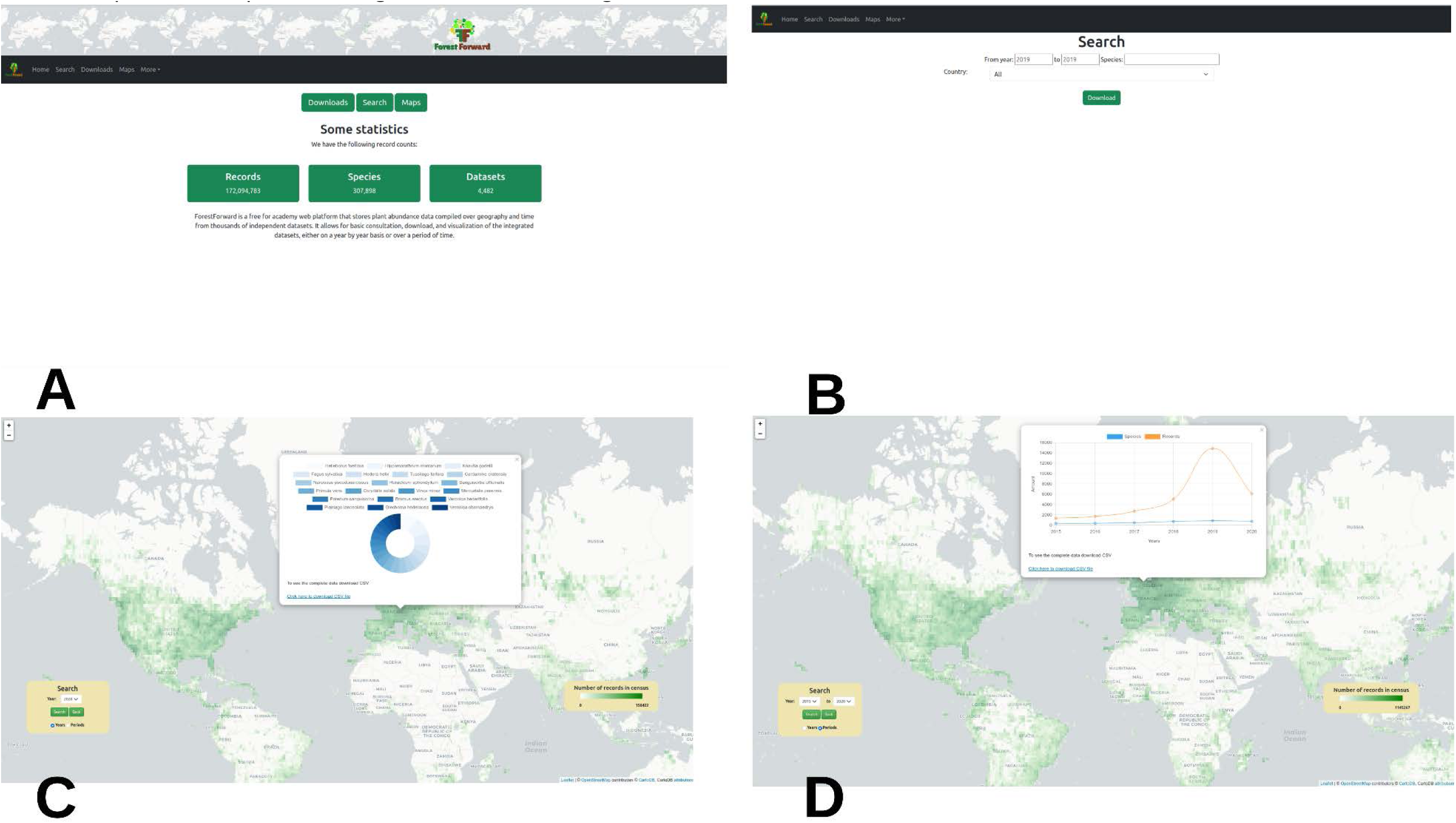
The ForestForward web user interface. A -. ForestForward entry page. **B -** After accessing the ForestForward web server users can go to the search view of the data. **C -** They can also access the world map directly and choose to view the data for any given year at any geographic location. By clicking on the geographic polygon, a pie chart with information about the species is displayed. A link in the chart allows users to download a csv file with the chart data. **D -** In the map view users can also display how the number of censed plants and species changed over time, and again download the data in csv format.

Second, users can access the compiled data on a world map (Figure 4C and Figure 4D). Overlaying the map, there is a layer with a square grid, with a resolution of 1x1 degrees of longitude and latitude. ForestForward colors the grid according to the amount of data available for every square in that year or period. The color scale goes from dark green (highest number of records) to transparent (no records for that region over the relevant period). Users can customize this view. They can choose to represent any other year of data from 1970 to 2020, any period of five years, or any decades over the same range (Figure 4D). When the view of interest for any year is selected and loaded, users can click any position in the map, and a pie chart with the percentage of plant species in that grid is shown (Figure 4C). When the view of interest for any decade is selected and loaded, clicking a position on the map will display a time course of the number of trees and species in the geographical polygon (Figure 4D). A link in the map allows users to download a cvs (Comma Separated Values) file containing all the data used to create the charts.

### Access

GBIF data have an open access policy that implements the principles of FAIR (findable, accessible, interoperable and reusable) (35) and open science, allowing unrestricted access to all researchers.

ForestForward follows the same principles. Our platform allows open access to the data, and every grouped data has its own id number. The downloadable csv format files enable interoperability. Users can download the data by years, by geographic location, and directly from the maps.

## Discussion

Access to high-quality data is crucial for researchers, and open access is key to enabling seamless data usage. While many researchers collect their own datasets through fieldwork and experiments, they also rely on data shared by colleagues, governmental agencies, and intergovernmental organizations. However, access to these datasets is often restricted, limiting collaboration and data sharing within the broader scientific community.

Establishing an open access database that integrates data from multiple sources in a cohesive, well-structured manner is essential for advancing scientific research. A platform adhering to the principles of FAIR (Findable, Accessible, Interoperable, and Reusable) and open science would foster transparency and collaboration among researchers and institutions. Open access promotes knowledge exchange, facilitating interdisciplinary research and enabling the scientific community to address complex global challenges collectively.

Moreover, an integrated open access database enhances the reproducibility and replicability of scientific results. Researchers can access and verify data from various sources, allowing them to validate or build upon previous studies. This level of transparency strengthens scientific rigor, helping to identify potential errors or biases and contributing to the refinement of research outcomes.

By offering access to diverse datasets, such a database facilitates the exploration of new research avenues and the discovery of novel patterns. It encourages interdisciplinary collaboration and integration across different fields, leading to innovative insights and breakthrough discoveries. Additionally, a platform aligned with FAIR principles ensures equitable access to data, enabling researchers worldwide to contribute to and benefit from shared knowledge, fostering global scientific progress.

### Data Quality and Pre-filtering in ForestForward

Data Extract, Transform, and Load (ETL) process is a semi-automated approach used to maintain an up-to-date data warehouse. The Extract phase focuses on acquiring high-quality data from multiple sources. Identifying and utilizing as many open access data sources as possible is critical to ensure data integrity and integration. Regular communication with these sources is necessary to facilitate periodic data updates. The Transform phase involves processing large volumes of data to meet the requirements of the data warehouse, which demands significant time and computational power. Efficient data integration requires query optimization and parallel execution, maximizing server core usage. The use of indexing within the Plantae collection has further reduced query processing time by pre-ordering the data. The Load phase involves storing and processing the data to ensure continuous updating, availability, and accessibility. These combined efforts will support the development of a robust geographic and statistical querying tool. This initiative represents a collaborative effort by researchers dedicated to enhancing access to high-quality data for the broader community.

To prioritize data quality, a pre-filtering process was implemented before integrating datasets from the Global Biodiversity Information Facility (GBIF). This pre-filtering ensured that only data with well-defined geographic coordinates and species identification were extracted, significantly improving data completeness and accuracy. As a result, the data integrated into ForestForward are three times more complete than those reported by Serra-Diaz (36), where only 20% of the analyzed data were considered complete and high-quality. This rigorous data curation enhances the reliability of ForestForward for spatiotemporal biodiversity analyses.

It is important to note that ForestForward tracks species occurrences over time in different regions. However, changes in the number of occurrence records do not necessarily indicate actual variations in species abundance. These fluctuations may reflect differences in data collection efforts rather than true population changes. For instance, an increase in records may result from more frequent sampling rather than population growth, while a decrease may reflect reduced surveying rather than a population decline. This distinction must be considered when interpreting the data.

### Data Acquisition and Integration Across Platforms

Data acquisition methods vary among biodiversity platforms. Some, such as the Global Biodiversity Information Facility (GBIF), Atlas of Living Australia (ALA), Global Forest Biodiversity Initiative (GFBI), Global Root Traits (GRooT), and World Flora Online (WFO), allow direct data contributions from researchers. Others, like ForestForward, Global Naturalized Alien Flora (GloNAF), and the Global Inventory of Floras and Traits (GIFT), aggregate data from multiple sources, requiring careful handling of multisource datasets to ensure consistency and compatibility.

These platforms manage various types of data, including geospatial data, plant images, and taxonomic information (e.g., species, genus, family, phylum). Specialized platforms like GRooT, WFO, and GIFT focus on plant physiology and morphology. Table 1 outlines the characteristics of these applications.

**Table 1.** Features of available query tools.

| Feature | Forest<br>Forward | GBIF | GFBI | ALA | GRooT | GloNAF | WFO | GIFT |
| --- | --- | --- | --- | --- | --- | --- | --- | --- |
| **Data acquisition & Integration** |  |  |  |  |  |  |  |  |
| Individual Contributors | No | Yes | Yes | Yes | Yes | No | Yes | No |
| Unique data structure | Yes | No | Yes | No | Yes | Yes | Yes | Yes |
| Integrates datasets | Yes | Yes | Yes | Yes | Yes | Yes | Yes | Yes |
| **Data types** |  |  |  |  |  |  |  |  |
| GIS | Yes | Yes | Yes |  | No | Yes | No | Yes |
| Plant images | No | Yes | No | Yes | No | No | Yes | No |
| Plant Species | Yes | Yes | Yes | Yes | Yes | Yes | Yes | Yes |
| Plant Genus | Yes | Yes | Yes | Yes | Yes | Yes | Yes | Yes |
| Plant Families | No | Yes | Yes | Yes | No | Yes | Yes | No |
| Plant Phylum | Yes | Yes | Yes | Yes | No | Yes | Yes | No |
| Physiology | No | No | No | No | Yes | No | No | Yes |
| Morphology | No | No | No | No | Yes | No | Yes | Yes |
| **Searchable data fields** |  |  |  |  |  |  |  |  |
| GIS | Yes | Yes | No | Yes | No | Yes | No | Yes |
| Species | Yes | Yes | No | Yes | No | Yes | Yes | Yes |
| Year | Yes | Yes | No | Yes | No | No | No | No |
| **Data Access** |  |  |  |  |  |  |  |  |
| Free | Yes | Yes | No | Yes | Yes | Yes | Yes | Yes |
| **Download** |  |  |  |  |  |  |  |  |
| Data availability for download | Yes | Yes | No | Yes | Yes | Yes | Yes | Yes |
| **Data analysis and representation** |  |  |  |  |  |  |  |  |
| Descriptive statistics | Yes | Yes | No | Yes | No | No | No | No |
| Graphical representation of statistics | Yes | No | No | No | No | No | No | No |

Search capabilities also differ across platforms. ForestForward, for instance, supports geospatial queries, allowing users to search by geographical coordinates. Most platforms support taxonomy-based searches, particularly by species, while platforms like GBIF and ForestForward also allow time-based searches, enabling the analysis of species occurrences over specific time periods in given regions.

While most platforms adhere to the principles of open access, GFBI restricts data to consortium members who contribute their own datasets. Similarly, data download options vary, with some platforms requiring a free account for access.

### Analytical Capabilities

Analytical and visualization tools are critical for biodiversity research. For example, ForestForward offers geospatial representations, basic statistical analyses, and descriptive graphical presentations. Its interactive maps display tree species distributions, aiding the understanding of ecological patterns, biodiversity assessments, and conservation decision-making. Temporal analyses, which examine data changes over time, are particularly important for tracking species distributions, assessing climate change impacts, and studying human interventions on ecosystems (37, 38).

The ability to conduct temporal analyses of species occurrence records is essential for understanding biodiversity changes over time. Detecting patterns in ecosystem changes requires long-term, multisource data collection. ForestForward facilitates this by offering tools that visualize changes over time, a feature lacking in GBIF. These capabilities provide researchers, forest managers, and conservationists with valuable tools for data exploration, analysis, and decision-making.

ForestForward can be useful for researchers studying species distribution over time in specific regions, helping them obtain integrated data to identify patterns. The platform’s development ensures that the scientific community has free access to integrated forestry data.

### Future Improvements for ForestForward

Future development of ForestForward will focus on three key areas. First, the integration of additional data types, such as climate, biome, and land use data, will enrich biodiversity trend analyses over time. These datasets will enable researchers to examine how environmental changes affect species distributions and ecosystem dynamics. Second, an Application Programming Interface (API) will be developed to facilitate direct integration with other web applications. Finally, in the longer term, machine learning algorithms will be incorporated to support advanced data analysis, offering predictive models for biodiversity trends and helping to identify critical areas for conservation and management.

In conclusion, integrating high-quality, open-access data into platforms like ForestForward enhances our understanding of global biodiversity patterns and enables more effective monitoring of changes. By combining multisource datasets and improving analytical tools, researchers can better understand the effects of climate change and human activities on ecosystems, ultimately supporting informed conservation efforts.

## Acknowledgements

The authors are members of the consolidated research group 2021SGR1353, accredited by Generalitat de Catalunya.

## Author contributions

E.T. designed the study, created the database, designed and developed the application, deployed the website, and wrote the manuscript. J.M. and F.S. deployed and debugged the website, revised the manuscript. R.A. designed the study, created the database and application, wrote the manuscript, coordinated the work.

---

**Supplementary Figure S1.**
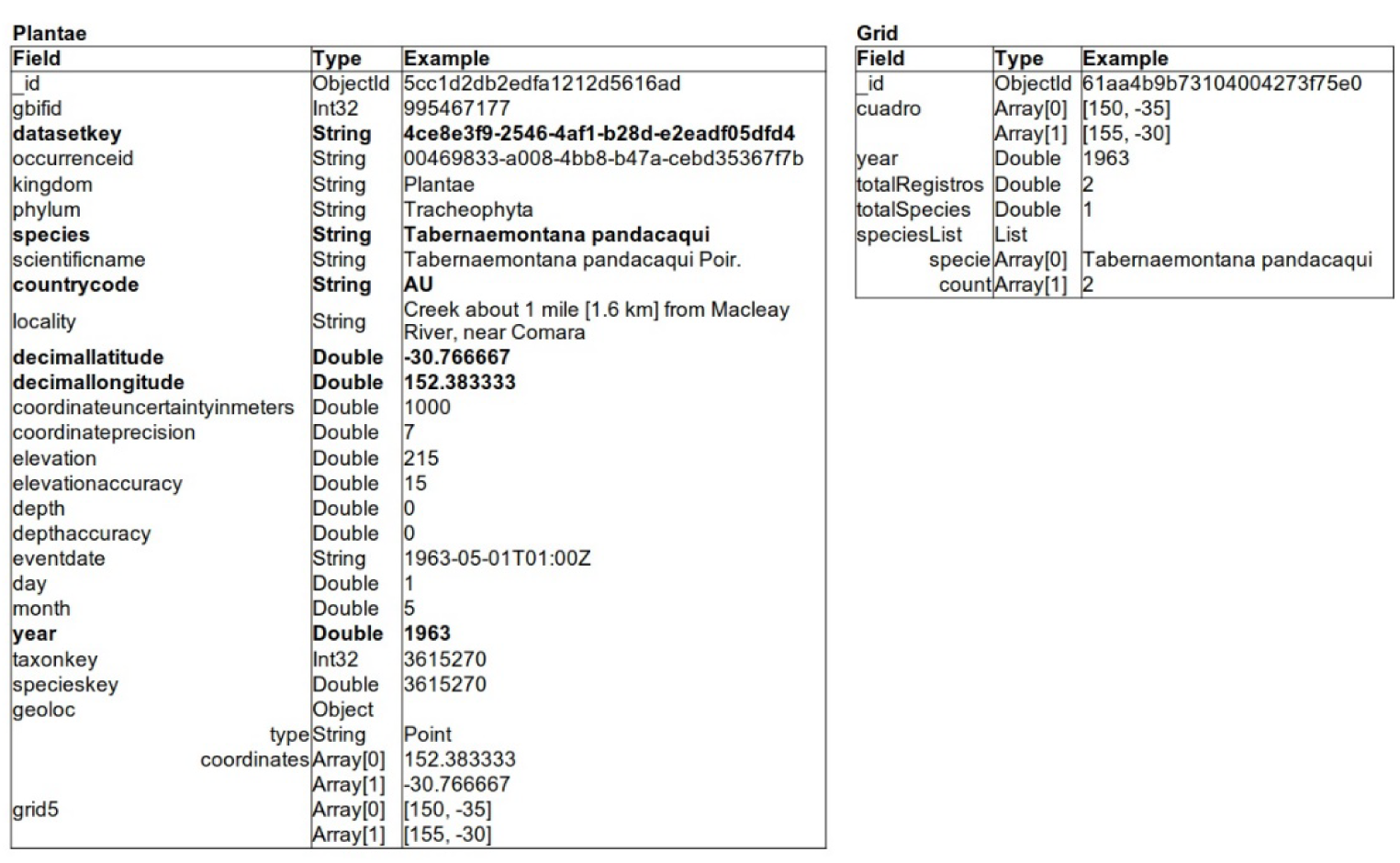
Schematic database structure for ForestForward.

